# LieOTMap: A Differentiable Approach to Cryo-EM Fitting via Lie-Theoretic Optimal Transport

**DOI:** 10.1101/2025.09.16.676533

**Authors:** Yue Hu, Zanxia Cao, Yingchao Liu

**Affiliations:** School of Bioengineering, Qilu University of Technology (Shandong Academy of Sciences), No. 3501 Daxue Road, Jinan, Shandong, China; Shandong Provincial Key Laboratory of Biophysics, Institute of Biophysics, Dezhou University, Dezhou 253023, China; Shandong Provincial Hospital, Shandong First Medical University

## Abstract

**Motivation:** The integration of high-resolution atomic models with lower-resolution cryo-electron microscopy (cryo-EM) maps is a fundamental task in structural biology. However, this rigid-body fitting problem is challenged by complex scoring landscapes and the non-differentiable nature of standard structural similarity metrics like TM-score, precluding their direct use in modern gradient-based optimization pipelines.

**Method:** We present LieOTMap, a novel, fully differentiable framework for cryo-EM fitting. Our approach introduces two key innovations. First, we parameterize rigid-body transformations on the Lie algebra *se*(3), which provides a minimal, singularity-free representation of the motion. Second, we formulate a loss function based on the Sinkhorn divergence, a differentiable proxy for the Optimal Transport (Wasserstein) distance. This loss function serves as a robust, geometrically meaningful surrogate for non-differentiable scores, allowing us to leverage powerful gradient descent optimizers to navigate the search space.

**Results:** We demonstrate the effectiveness of LieOTMap by fitting the apo-state structure of *E. coli* GroEL (PDB: 1aon) into the ATP-bound state cryo-EM map (EMD-1046). Our method successfully navigates a large-scale conformational change, achieving a highly accurate final RMSD of 3.08 Å with respect to the ground-truth structure (PDB: 1GRU). This result showcases the power of combining Lie-theoretic representations with differentiable geometric loss functions for complex structural alignment tasks.

**Availability:** The source code is available at https://github.com/YueHuLab/LieOTMap.

## 1 Introduction

Hybrid methods that integrate data from multiple structural biology techniques have become indispensable for elucidating the architecture of complex macromolecular assemblies. A prominent example is the fitting of high-resolution atomic models, typically obtained from X-ray crystallography or NMR, into lower-resolution density maps generated by cryo-electron microscopy (cryo-EM) (Ranson et al., 2001). This process is crucial for interpreting the cryo-EM map in atomic detail, understanding conformational changes, and formulating functional hypotheses.

The core of the fitting problem is to determine the precise rigid-body transformation (rotation and translation) that best aligns the atomic model with the density map. A variety of algorithms have been developed to address this challenge. Traditional methods often rely on maximizing a cross-correlation score, but their scoring landscapes can be complex and difficult to optimize. More recent approaches frame the task as a point cloud registration problem.

However, a key limitation of many structural biology tools is their reliance on metrics, such as the popular TM-score (Zhang and Skolnick, 2005), that are not differentiable. This prevents the direct use of highly effective gradient-based optimization algorithms that are at the heart of modern machine learning. Furthermore, the parameterization of 3D rotations itself can introduce challenges, with representations like Euler angles suffering from singularities (gimbal lock).

In this work, we introduce a novel framework, LieOTMap, that is differentiable end-to-end. We address the aforementioned challenges by integrating two powerful mathematical concepts. First, we parameterize rigid motion using the Lie algebra *se*(3), a minimal, 6-parameter representation that is continuous and free of singularities (Murray et al., 1994). Second, we employ the Sinkhorn divergence, a differentiable and computationally efficient approximation of the Wasserstein distance from Optimal Transport (OT) theory (Cuturi, 2013), as a robust geometric loss function. This serves as a differentiable proxy for the non-differentiable structural similarity scores, enabling a complete gradient-based optimization pipeline.

We demonstrate the efficacy of our method on the conformational change of the *E. coli* GroEL/GroES chaperonin complex, a challenging and biologically significant test case. Our method successfully rediscovers the correct orientation, achieving a final RMSD of 3.08 Å with respect to the gold-standard structure.

## 2 Methods

### 2.1 Point Cloud Representation

Let the mobile atomic structure be represented by a point cloud 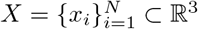, where each *x*_*i*_ is the coordinate of a C-alpha atom. The target cryo-EM density map, *ρ*(*v*), is a scalar field on a 3D grid. We convert this map into a target point cloud, 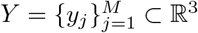, by applying a sigma-level threshold. First, the mean (*µ*_+_) and standard deviation (*σ*_+_) of all positive density values are computed. A threshold *T* = *µ*_+_ + *k* · *σ*_+_ is defined (in our work, *k* = 3.0). The target point cloud *Y* consists of the real-space coordinates of all voxels *v*_*j*_ where *ρ*(*v*_*j*_) *> T* .

### 2.2 Rigid Motion on the Lie Algebra

A rigid-body transformation is an element of the Special Euclidean group, SE(3). To facilitate unconstrained optimization, we use the Lie algebra *se*(3), which is the vector space tangent to the SE(3) manifold at the identity. An element of *se*(3) is a 6-dimensional vector *ξ* = (*v, ω*) ∈ ℝ^6^, called a “twist”, which can be mapped to a 4 × 4 matrix representation 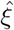. The transformation matrix *T* ∈ SE(3) is recovered via the matrix exponential map, 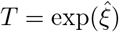 (Murray et al., 1994). This allows us to parameterize the entire rigid transformation with just six unconstrained variables, which are the parameters we optimize.

### 2.3 Differentiable Loss: Sinkhorn Divergence as a Proxy for TM-score

The de facto standard for evaluating protein structure similarity is the TM-score, which is defined as:

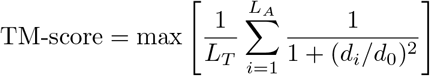

where *L*_*T*_ is the length of the target protein, *L*_*A*_ is the number of aligned residue pairs, *d*_*i*_ is the distance of the *i*-th pair, and *d*_0_ is a distance threshold. A key feature of the TM-score is its non-differentiability, arising from the maximization operation over all possible alignments. This prevents its direct use in a gradient-based search.

Our approach is to replace this non-differentiable metric with a differentiable surrogate that still captures the geometric essence of the alignment problem. We use the Sinkhorn divergence, *S*_*ϵ*_, which is derived from Optimal Transport theory (Cuturi, 2013). The regularized OT cost is:

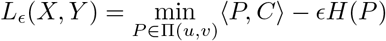

where *C*_*ij*_ = ∥*x*_*i*_ − *y*_*j*_∥ ^2^ is the cost matrix and *H*(*P*) is the entropy of the transport plan *P* . The Sinkhorn divergence is then defined as:

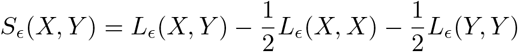

This function is fully differentiable with respect to the point coordinates *X* and provides a robust measure of dissimilarity between the two point clouds, serving as an excellent proxy objective for the true alignment quality.

### 2.4 Optimization

By combining these concepts, our final optimization problem is to find the optimal twist coordinates *ξ*^*^:

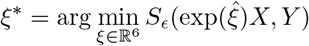

We solve this unconstrained optimization problem using the Adam optimizer. The gradients of the loss function with respect to the 6 parameters of *ξ* are computed via automatic differentiation, and the parameters are updated iteratively.

## 3 Results and Discussion

We demonstrated our method on the task of fitting the chaperonin GroEL. We used the crystal structure of GroEL in its apo state (PDB: 1aon) as our mobile model and aimed to fit it into the 3D cryo-EM density map of the ATP-bound GroEL/GroES complex (EMD-1046) (Ranson et al., 2001). The accepted solution for this map is the refined gold-standard structure (PDB: 1GRU), which we used to validate our final result.

The optimization was performed for 1000 steps. The algorithm showed rapid convergence, with the RMSD relative to the gold standard dropping from an initial value of 15.86 Å to below 3.1 Å in the first 800 steps. The final RMSD after 1000 steps was 3.08 Å, indicating a highly accurate fit. The key metrics of the fitting process are summarized in Table 1.

**Table 1:** Summary of the fitting results for PDB 1aon into map EMD-1046.

| Stage | Metric | Value |
| --- | --- | --- |
| Initial Alignment (Post-CoM) | RMSD | 15.86 Å |
| Optimization (500 steps) | RMSD | 3.59 Å |
| Optimization (Final) | RMSD | 3.08 Å |
| Optimization (Final) | TM-score | 0.993847 |

The visual results of this process are depicted in Figure 1. Panel A shows the target point cloud derived from the density map. Panel B shows the final superposition of our fitted model (1aon, cyan) and the gold-standard structure (1GRU, magenta). The excellent visual overlap confirms the low RMSD value and the success of our fitting procedure.

**Figure 1.**
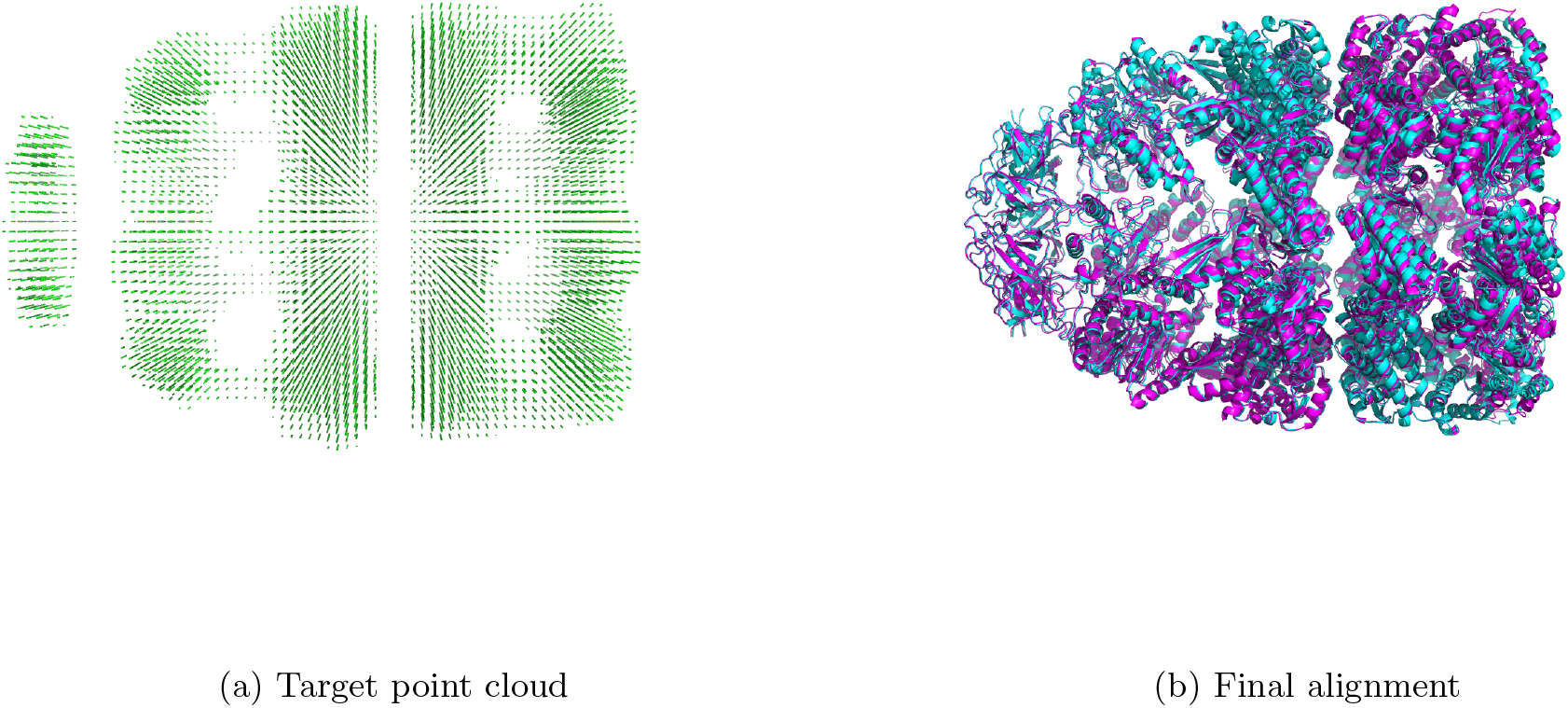
Visual summary of the fitting results. (A) The target point cloud (*Y*) generated from the EMD-1046 density map using a 3.0-sigma threshold. (B) Superposition of our final fitted model (1aon, colored cyan) and the gold-standard structure (1GRU, colored magenta). The close overlap demonstrates the high accuracy of the fit.

The successful fitting of 1aon into the EMD-1046 map, a non-trivial case involving significant conformational differences, demonstrates the power of our approach. The ability to achieve a 3.08 Å RMSD confirms that our method can accurately determine the rigid-body component of complex conformational changes. The use of a Lie algebra representation for transformations provides a minimal, unconstrained parameter space for gradient-based optimization, elegantly avoiding the pitfalls of other rotation representations. The Sinkhorn divergence, as a loss function, is well-suited for comparing point clouds of different sizes and is robust to noise, a common feature of cryo-EM data.

## 4 Conclusion

We introduced a novel, fully differentiable method for fitting atomic structures into cryo-EM maps by combining a Lie-theoretic representation of rigid motion with an Optimal Transport-based loss function. This approach allows for efficient and robust gradient-based optimization. We validated its high accuracy on a real-world biological system, demonstrating its potential as a powerful tool for hybrid structural modeling.

## References

[1] Cuturi, M. (2013). Sinkhorn Distances: Lightspeed Computation of Optimal Transport. Advances in Neural Information Processing Systems, 26.

[2] Murray, R. M., Li, Z., Sastry, S. S. (1994). A mathematical introduction to robotic manipulation. CRC press.

[3] Ranson, N. A., Farr, G. W., Roseman, A. M., Gowen, B., Fenton, W. A., Horwich, A. L., Saibil, H. R. (2001). ATP-bound states of GroEL captured by cryo-electron microscopy. Cell, 107(7), 869–879.

[4] Zhang, Y., Skolnick, J. (2005). TM-align: a protein structure alignment algorithm based on the TM-score. Nucleic acids research, 33(7), 2302–2309.

